# Circadian Rhythm Disruption Results in Visual Dysfunction

**DOI:** 10.1101/2020.12.14.422683

**Authors:** Deepa Mathew, Ashay D. Bhatwadekar

## Abstract

Circadian rhythm disruption (CRD) contributes to the development of multiple metabolic and neurodegenerative diseases. However, its effect on vision is not understood. We evaluated the impact of CRD on retinal morphology, physiology, and vision after housing mice in a disruption inducing shorter light/dark cycle (L10:D10). Interestingly, the mice under L10:D10 exhibited three different entrainment behaviors; ‘entrained’, ‘freerunning’, and ‘zigzagging.’ These behavior groups under CRD exhibited reduced visual acuity, retinal thinning, and a decrease in the number of rod photoreceptors. Intriguingly, the electroretinogram response was decreased only in the mice exhibiting ‘entrained’ behavior. The retinal proteome showed distinct changes with each entrainment behavior. These results demonstrate that CRD leads to photoreceptor degeneration and visual dysfunction. We uniquely show the effect of entrainment behavior on retinal protein composition and physiology. Our data has broader implications in understanding and mitigating the effect of CRD on vision health.

**Significance Statement:** Artificial light is increasingly in use for the past 70 years. The aberrant light exposure and round-the-clock nature of work lead to the disruption of a biological clock. The mammalian retina possesses an autonomous clock that could reset with light exposure. Yet, the effect of altered light exposure on vision is not studied. Here, we report visual dysfunction after exposure to a circadian disrupting altered light/dark cycle. We also report behavior dependent changes in retinal protein composition and function. Our data provide evidence for the negative impact of CRD on vision and its potential role in retinal diseases’ etiology.

## Introduction

Circadian rhythms are the near 24-hour patterns in physiology and behavior that persist in the absence of external stimuli. In mammals, a neuronal network in the suprachiasmatic nucleus (SCN) generates endogenous circadian rhythms and orchestrates the complex system of circadian oscillators throughout the body via behavioral, hormonal, and neuronal signals (1). The SCN maintains its phase coherence with the environmental light-dark cycles through neuronal projections from the retina via the retina-hypothalamic tract (1). Within the cells of SCN and every other tissue, circadian oscillation is driven by a cell-autonomous autoregulatory feedback loop of clock genes (2). Environmental and systemic inputs could feed into this core clock mechanism (2), possibly modulating its precision and flexibility while adapting to perturbations.

Disruption of this fine-tuned machinery is widespread in the modern world with the extensive use of artificial lighting, round the clock nature of work, and trans meridian travel. Chronic circadian rhythm disruption (CRD) has been identified as a risk factor for multiple metabolic, cardiovascular and neurodegenerative diseases (3, 4). Animal models with clock gene mutations also showed metabolic and neurodegenerative disorders (3). Moreover, exposure to shorter light-dark cycles resulted in metabolic and neuropsychiatric alterations in genetically intact mice (5, 6).

The mammalian retina possesses an autonomous circadian clock that can be entrained by light (7, 8), as well as the retinal clock is also exposed to systemic circadian time cues. Thus, retinal gene expression and function are tightly controlled by the circadian clock and exposure to light (7, 9). Moreover, ablation of clock gene *Bmal* in mice resulted in alterations of ocular gene expression, reduced electroretinogram (ERG) amplitude, and cone photoreceptor viability (9, 10). Mutant mice for BMAL or PER2 exhibited changes in their retinal vasculature, suggesting the role of clock genes in the pathogenesis of vascular eye diseases such as diabetic retinopathy (11, 12). However, the effect of altered light exposure on the visual system has not been evaluated. In this study, we exposed genetically intact mice to a shorter light-dark cycle (L10:D10) to assess the effect of CRD on different attributes of vision such as visual acuity, and retinal structure, function, and proteome.

## Results

### CRD resulted in different entrainment behaviors

C57BL /6J mice exposed to L10:D10 exhibited three distinct entrainment behaviors in their wheel-running activity: (**i**) ‘Entrained’ (E), (**ii**) ‘freerunning’ (F) and (**iii**) ‘zigzagging’ (Z). Mice that ‘entrained’ to L10:D10 restricted their activity only in the dark phase with a period of ~20 hours and a phase of entrainment of ≤1 hr to lights off (Fig.1B, 1E; Fig. S1). The mice which exhibited ‘freerunning’ did not entrain to the external light / dark cycle and showed a period of 23.09 ± 0.24 hours (Fig.1C, 1E; Fig. S2). The ‘zigzagging’ mice lacked a stable phase of entrainment and switched their period every 3-5 days between 21.48 ± 0.12 hours and 18.47± 0.21 hours (Fig. 1 D, 1E; Fig. S3). Lomb-Scargle periodogram analysis revealed a 20 hour and a >20 hour periodicities for ‘freerunning’ mice (Fig. S2), and a 20 hour period for ‘zigzagging’ mice (Fig. S3). Since these periods did not match the period calculated by fitting regression lines on activity onsets, we report the latter as well (Fig 1E). Three of the F mice initially showed unstable periodicities of 22.31±0.26, which shortened when light exposure was at the end of the active phase and lengthened when light exposure was at the beginning of the active phase (Fig. S2 D, E, H). These 3 mice settled at stable longer periodicities between 8-12 weeks. These distinct entrainment behaviors observed in L10:D10 were not tested under constant conditions.

**Fig 1.**
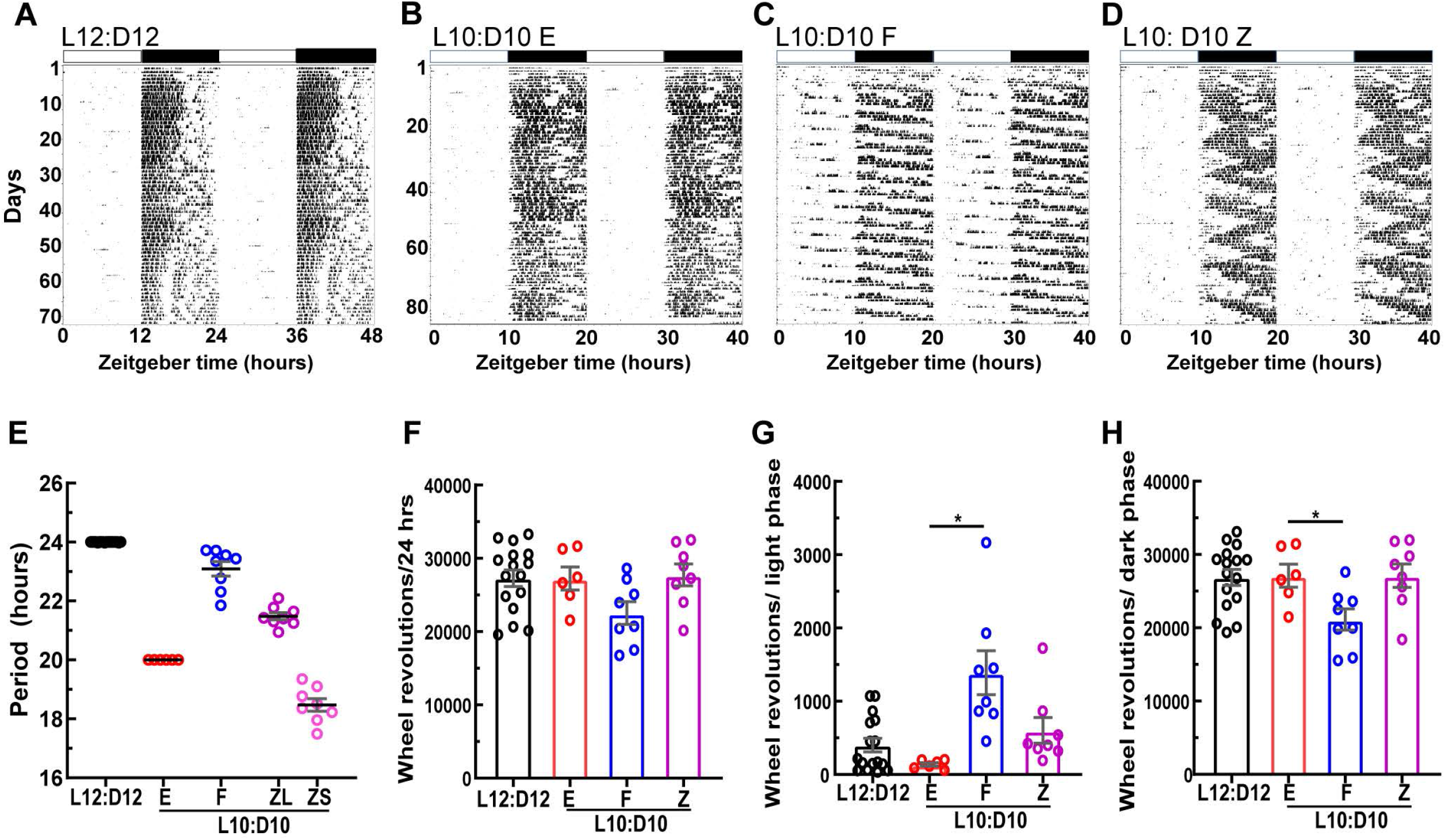
Mice under L10: D10 conditions display different entrainment behaviors. Representative double plotted actograms of wheel-running activity under control and L10:D10 conditions showing different behavior patterns; control (A), ‘entrained’ (B), ‘freerunning’ (C) and ‘zigzagging’ (D). Periods exhibited in each entrainment behavior, obtained by fitting regression lines on activity onsets on the actograms using Clocklab (E). Quantification of wheel-running activity (F) and its distribution to light (G) and dark phases(H). Shown are mean + SEM. * P ≤ 0.05 (Dunnet’s test).

Next, we quantified wheel revolutions to evaluate if it has any protective effect on retinal physiology(13). Total wheel-running activity was not significantly different between control and L10:D10 behavior groups (Fig.1F; W_3,15_=2.45, p=0.10). However, wheel revolutions differed significantly between behaviors when separated as light phase (Fig.1G; W_3,15_=9.24, p=0.001) and dark phase (Fig.1H; W_3,15_=3.83, p=0.03). The ‘freerunning’ mice had higher light phase activity (Fig.1G, p=0.02) and lower dark phase activity (Fig. 1H, p=0.04) compared to ‘entrained’, since their activity was distributed across all phases of the L10:D10 light/dark cycle.

### CRD resulted in reduced visual acuity

To determine whether a visual function was affected by CRD, optomotor response tracking was performed, and visual acuity defined by spatial frequency threshold was measured. L10:D10 groups demonstrated significantly reduced visual acuity irrespective of their entrainment behavior (Fig. 2; W_3,30_=39.15, p<0.0001).

**Fig 2.**
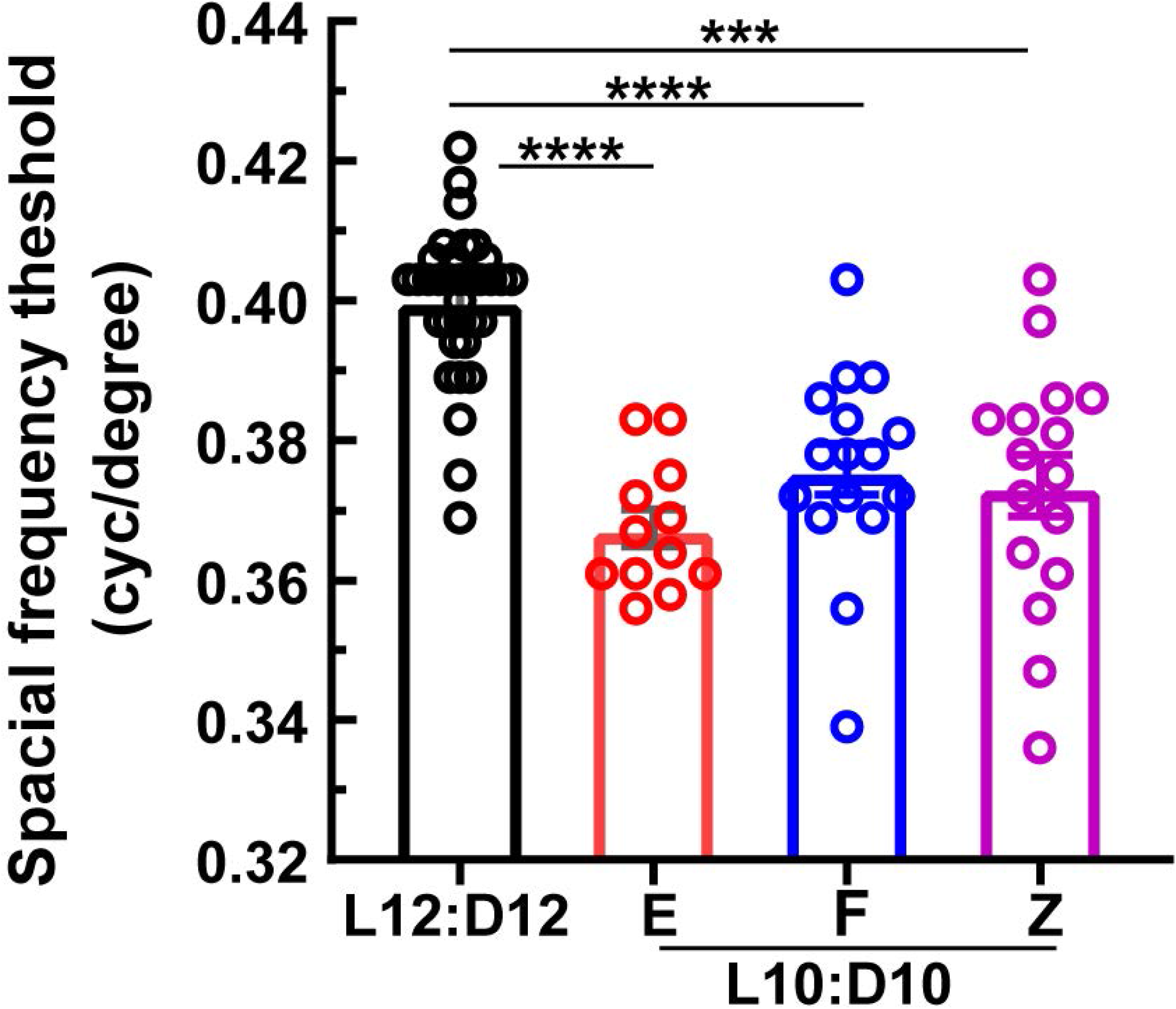
Decrease in visual acuity under L10:D10 conditions: L10:D10 mice showed reduced visual acuity when measured using an optomotor response tracking system. Shown are mean + SEM. *** P ≤ 0.001, **** P ≤ 0.001 (Dunnet’s test).

### CRD results in reduced retinal thickness and loss of photoreceptor cells

To investigate the effect of CRD on retinal structure, spectral-domain optical coherence tomography (SD-OCT) and immunohistochemistry were performed. The overall retinal thickness measured from SD-OCT b-scans was significantly reduced in all L10:D10 behavior groups (Fig. 3A, B; W_3,23.72_=15.35, p<0.0001). Immunohistochemistry using rhodopsin and cone arrestin antibodies was performed to quantify photoreceptors after CRD (Fig. 4A.). The number of rod photoreceptors was reduced in all L10:D10 behavior groups (Fig. 4B; W_3,6.1_=11.75, p=0.006). While the cone photoreceptors tended to be reduced in all L10: D10 groups, the difference was statistically significant only in the ‘entrained’ mice (Fig. 4C; W_3,5.54_=6.25, p=0.032).

**Fig 3.**
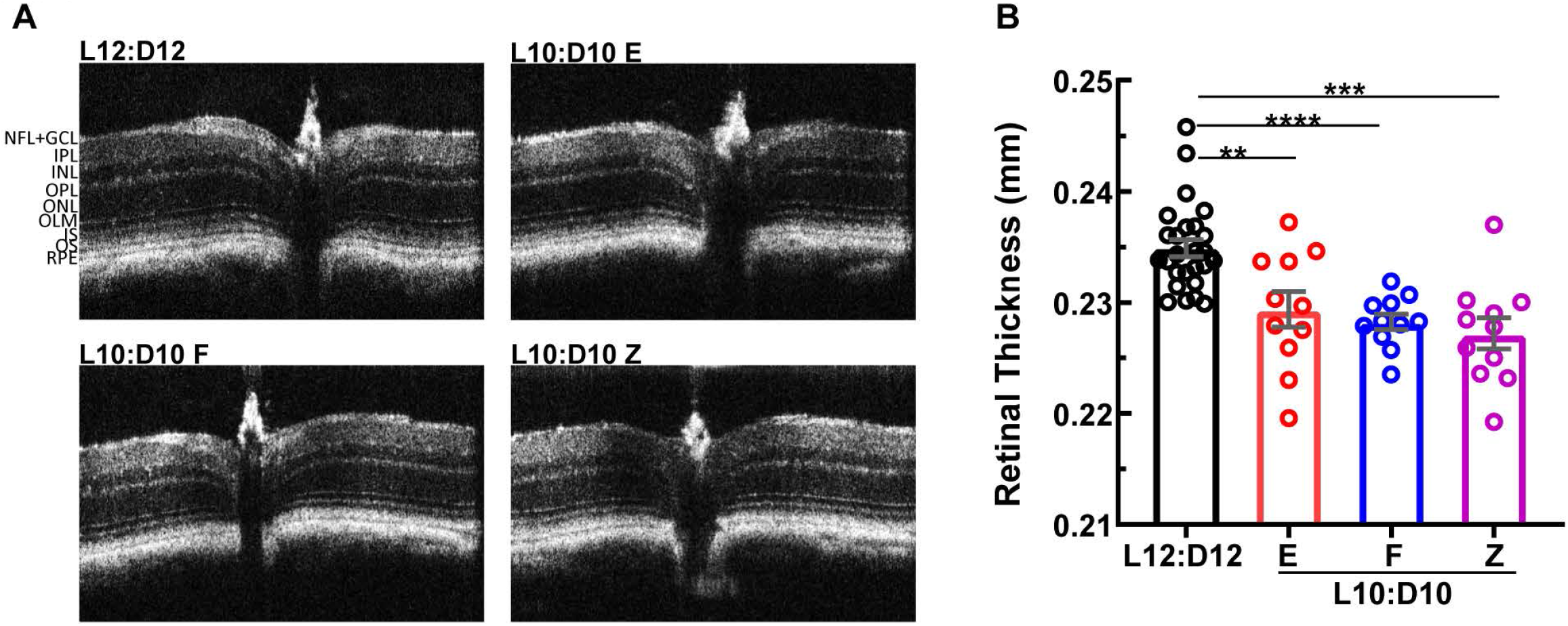
Retinal thinning under L10:D10 conditions. Representative Spectral Domain-Optical Coherence Tomography (SD-OCT) B-scans of the mouse retina (A), showing central optic nerve head and retinal sublayers. L10:D10 mice showed reduced total retinal thickness(B). Shown are mean + SEM. ** P ≤ 0.01, *** P ≤ 0.001, **** P ≤ 0.001 (Dunnet’s test).

**Fig 4.**
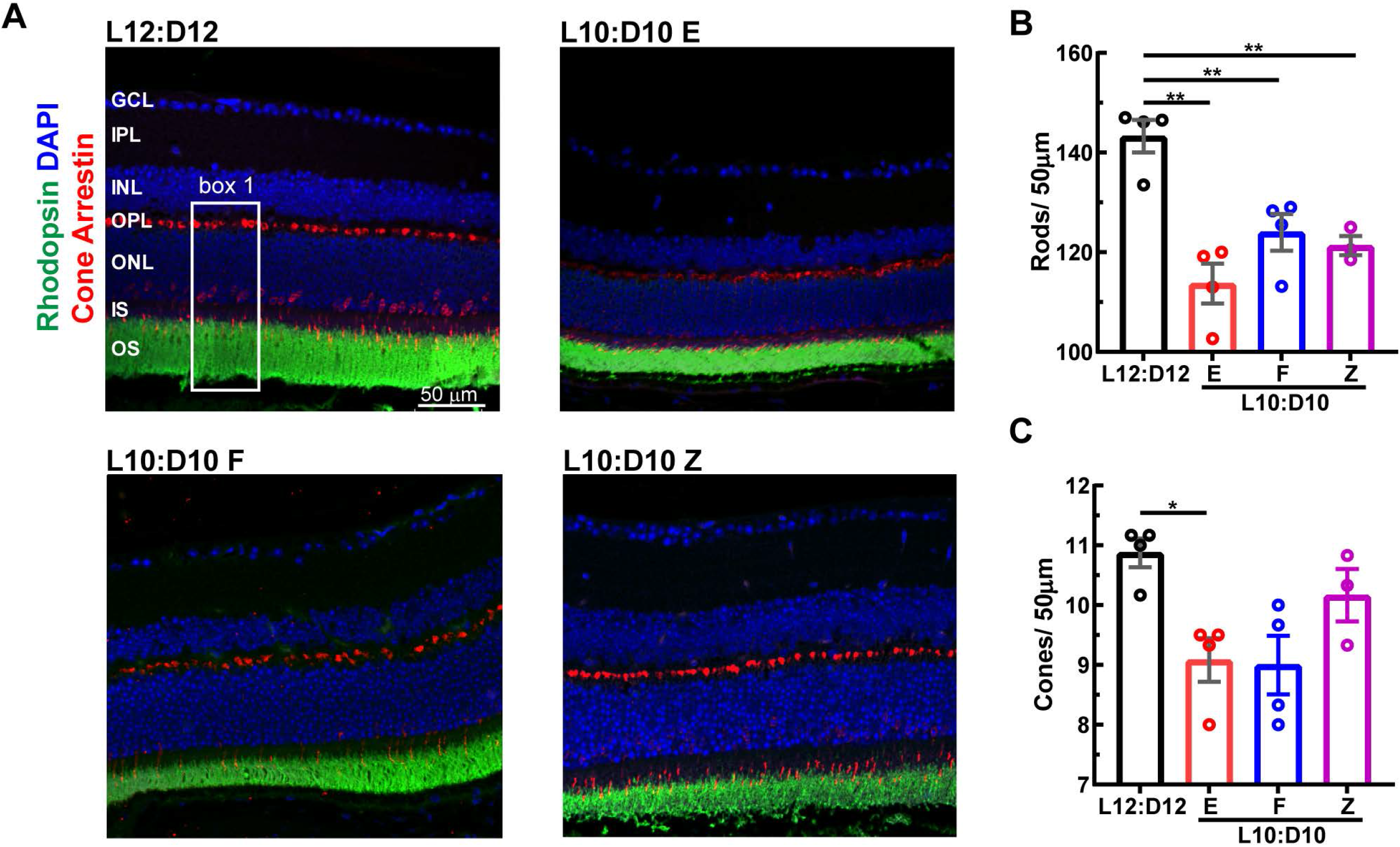
A decrease in the number of photoreceptor cells in L10:D10 treatment. Retinal sections were immunostained for photoreceptors (A). Cell densities of rods (total nuclei – cones, in box1) (B) and cones (C) per 50µm. Shown are mean + SEM. * P ≤ 0.05, ** P ≤ 0.01 (Dunnet’s test).

### Scotopic electroretinogram a-wave and b-wave amplitudes were reduced after CRD depending on the entrainment behavior

To assess retinal function, ERG response to flashes of light, under scotopic (rod driven) and photopic (cone isolated) conditions were recorded. Under scotopic conditions L10:D10 entrainment behavior showed a significant effect on the amplitudes of a-wave (Fig. 5A; flash intensity- F_2,216_=312, p<0.0001, behavior- F_3,216_=9.29, p<0.0001, interaction- F_3,216_=9.29, p=0.34), b-wave (Fig. 5B; flash intensity- F_2,216_=9.8, p<0.0001, behavior- F_6,216_=24, p<0.0001, interaction- F_6,216_=0.31, p=0.93) and average oscillatory potential (Fig. 5C; flash intensity- F_2,216_=39.2, p<0.0001, behavior- F_3,216_=14.7, p<0.0001, interaction- F_6,216_=1.3, p=0.27). Both a-wave that originates from photoreceptor cells and b-wave that originates from bipolar cells were significantly reduced in the ‘entrained’ mice while those of ‘free running’ and ‘zigzagging’ mice were unaffected (Fig. 5A, B). L10:D10 entrainment behavior showed effect on scotopic a-wave peak latency (Fig. S4A; flash intensity- F_2,216_=335, p<0.0001, behavior- F_3,216_=2.86, p=0.04, interaction- F_6,216_=0.43, p=0.86) while it did not show any effect on b-wave peak latency (Fig. S4B; flash intensity- F_2,216_=384, p<0.0001, behavior- F_3,216_=2.58, p=0.06, interaction- F_6,216_=0.76, p=0.6). These results further emphasize the loss of rod photoreceptor function in the ‘entrained’ mice

**Fig 5.**
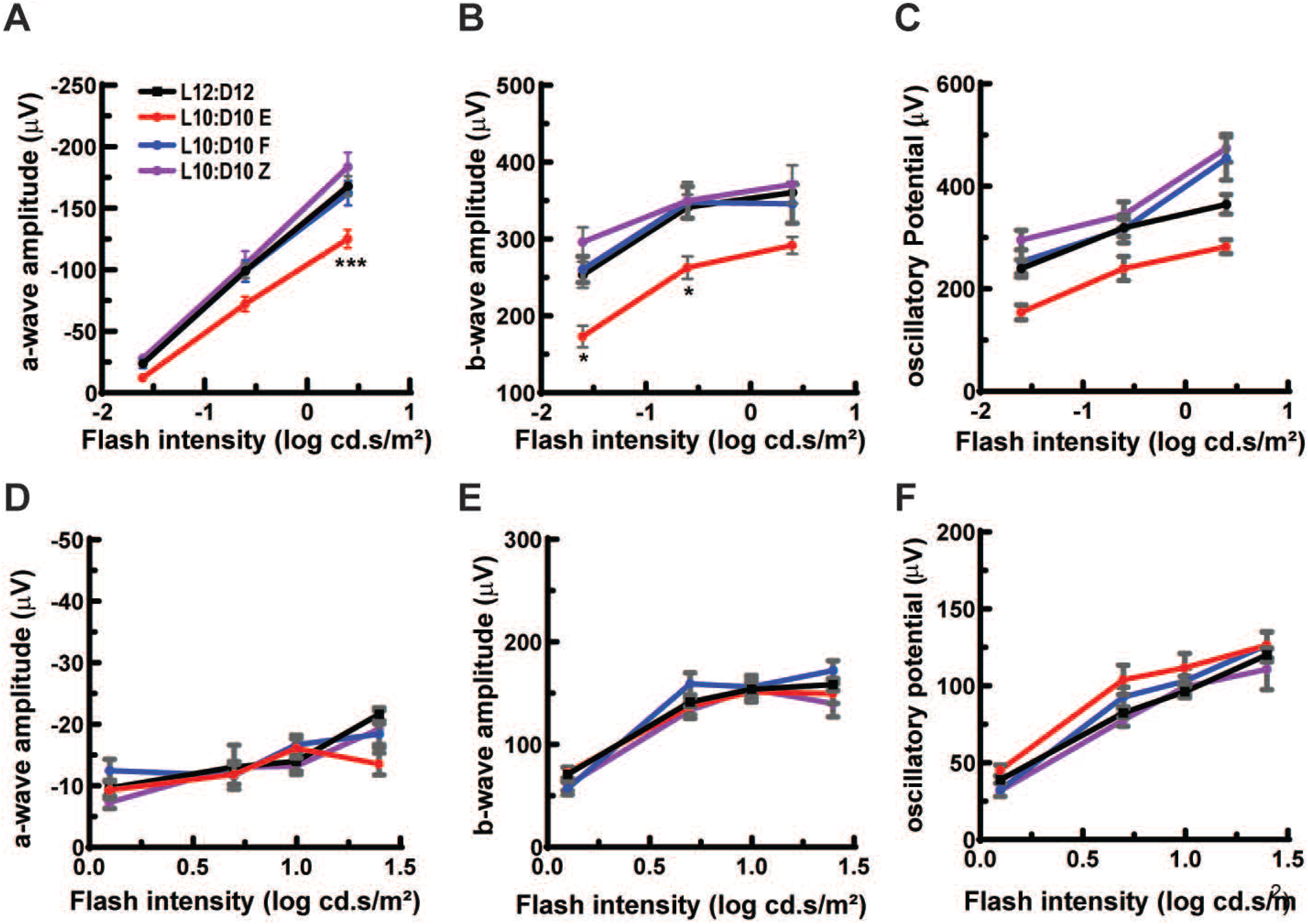
Altered ERG responses in mice under L10: D10 conditions. Entrained mice showed reduced amplitudes for scotopic a-wave (A) and b-wave (B) and average oscillatory potential (C), while ‘freerunning’ and ‘zigzagging’ did not show any difference. Amplitudes of photopic a-wave (D) and b-wave (E) and average oscillatory potential (F) were not significantly different between the groups. N=17 (L12:D12), 6 (L10:D10 E), 8 (L10:D10 F), 8(L10:D10 Z). Shown are mean + SEM. * P ≤ 0.05, *** P ≤ 0.001 (Tukey’s test).

Under photopic conditions a-wave and b-wave amplitudes were not significantly different between control and L10:D10 behavior groups (Fig. 5D; flash intensity- F_3,288_=17.4, p<0.0001, behavior- F_3,288_=1.35, p=0.26, interaction- F_9,288_=1.71, p=0.09, Fig. 5E; flash intensity- F_3,288_=111.9, p<0.0001, behavior- F_3,288_=2.05, p=0.11, interaction- F_9,288_=1.14, p=0.34). Entrainment behavior did show an effect on photopic oscillatory potential amplitude, a-wave peak latency and b- wave peak latency (Fig.5F; flash intensity- F_3,275_=138, p<0.0001, behavior- F_3,275_=4.47, p=0.004, interaction- F_9,275_=0.74, p=0.09, Fig. S6A; flash intensity- F_3,288_=20.7, p<0.0001, behavior- F_3,288_=4.4, p=0.005, interaction- F_9,288_=2.1, p=0.03, S4B; flash intensity- F_3,288_=6.2, p=0.0004, behavior- F_3,288_=9.8, p<0.0001, interaction- F_9,288_=4.2, p<0.0001). The free running group showed significantly shortened peak latencies for a- wave and b-wave at a flash intensity of 1.25 cd.s/m^2^(Fig. S4 A, B).

### CRD resulted in distinct changes in the retinal proteome with each entrainment behavior

To identify the molecular alterations underlying the observed changes in retinal physiology, we performed a proteomic analysis of the retina from all entrainment behaviors. Overall a total of 7620 proteins were identified. Interestingly, the quantitation of protein abundance revealed distinct proteomic changes in the retina with each entrainment behavior, as demonstrated by the volcano plots (Fig. 6 A, C, E). The ‘entrained’ retina showed 317 proteins upregulated and 174 proteins downregulated. The ‘freerunning’ retina showed 149 proteins upregulated and 156 proteins downregulated. Surprisingly, the most significant number of proteins were differentially expressed in the ‘zigzagging’ behavior; 441 upregulated and 538 downregulated. A complete list of differentially expressed proteins is provided as supplementary information.

**Fig 6.**
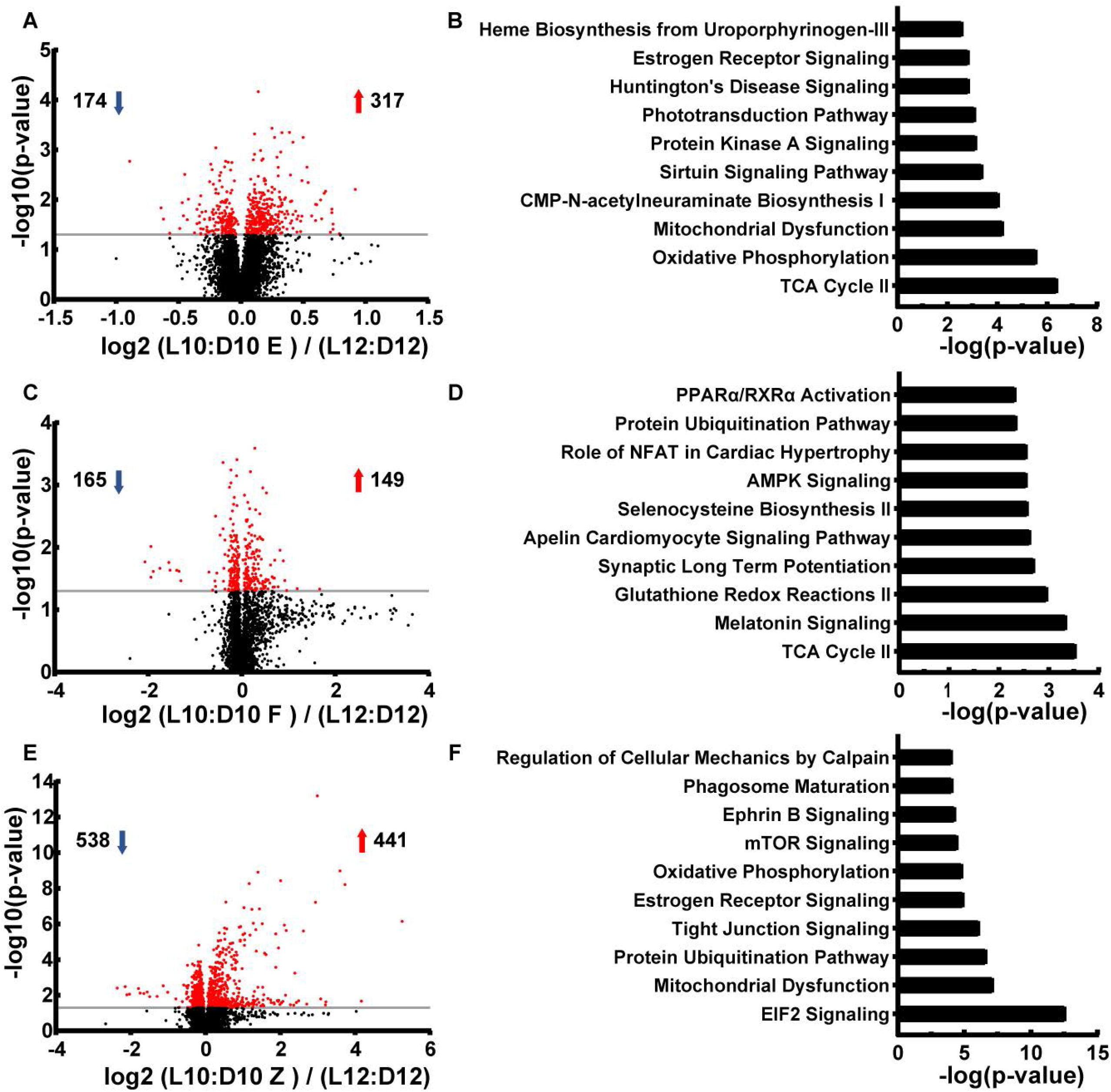
Distinct retinal proteomic changes and their associated pathways (IPA) in different L10:D10 entrainment behavior. Volcano plots demonstrating differentially regulated proteins in ‘entrained’ (A), ‘freerunning’ (C), and ‘zigzagging’ (E). Each data point represents a single quantified protein. The x-axis represents log fold change in abundance. The −log (p-value) is plotted on the y-axis. Proteins above the grey horizontal line indicate significance ≤ 0.05. Top 10 canonical pathways enriched from IPA analysis of differentially regulated proteins in Entrained (B), Freerunning (D), and zigzagging (F). The significance of functional enrichment is plotted on the x-axis as the −log (p-value) of Fisher’s exact test.

To gain more information about the molecular pathways associated with altered protein expression, we performed Ingenuity Pathway Analysis (IPA). This analysis predicted activation or inhibition of specific canonical pathways associated with each entrainment behavior (Fig.6 B, D, F; Supplementary table 1, 2, and 3). In the ‘entrained’ retina, pathways associated with mitochondrial function were affected. While, in the ‘freerunning’ group, multiple protective pathways were activated, including TCA cycle, long-term synaptic potentiation and AMPK signaling (14, 15). Melatonin signaling was also found to be activated in this behavior, suggesting alterations to their retinal clock. In the ‘zigzagging’ retina, ELF2 signaling, which was activated in endoplasmic reticulum stress, was found to be inhibited (16) while mTOR and Ephrin signaling were activated.

A comparative analysis of differentially regulated pathways revealed distinct changes between the behavior groups (Fig S5 A). The insulin secretion pathway was predicted as a top pathway differentially regulated between different groups, highly upregulated in entrained mice followed by free-running and zigzaggers. Interestingly TCA cycle pathway was predicted to be inhibited in the ‘entrained’ retina, while it was activated in the ‘free-running’ and ‘zigzagging’ retina, suggesting reduced retinal energy production after L10:D10 entrainment. Rho GDI signaling was significantly inhibited in the ‘entrained’ retina while it was activated in the ‘free-running’ and ‘zigzagging’ retina. A comparative analysis of predicted diseases and conditions of the brain and eye showed accelerated photoreceptor degeneration in the ‘entrained’ retina (Fig.S5 B).

## Discussion

The light exposure at the wrong time could result in CRD and, as such, be associated with metabolic and neurodegenerative diseases (4). Our study further demonstrates the negative impact of CRD on vision by showing a decrease in visual acuity, altered retinal structure as exhibited by a reduction in retinal thickness, photoreceptor degeneration and retinal function in ERG assessments. Changes in the above retinal parameters were coupled with the unique retinal proteome that mapped pathways integral to normal retinal function. Furthermore, we uniquely report different entrainment behaviors in C57BL/6J mice in response to a shorter light/dark cycle.

In our study, CRD led to reduced visual acuity, retinal thinning, and loss of rod photoreceptors irrespective of the entrainment behavior. Several possible mechanisms may explain why we see changes in retinal parameters due to a shorter light/dark cycle. First of all, our studies indicate that a reduced number of photoreceptors could have contributed to the observed retinal thinning and loss of visual acuity. Secondly, alterations in the neuronal connectivity upstream of bipolar cells either at the level of retinal ganglion cells or in their visual cortical circuitry could also decrease visual acuity. Previous studies support this assertion by showing a deceased complexity and loss of dendritic lengths in neurons in their prelimbic prefrontal cortex and decreased cognitive flexibility under similar light and dark conditions (5).

Our findings demonstrate that a decrease in rod-driven ERG function was only observed in the ‘entrained’ behavioral group, suggesting a more significant impact of CRD on this group’s retinal physiology. The proteome analysis provides some mechanistic explanation for this. Proteomic changes associated with the TCA cycle, oxidative phosphorylation, mitochondrial dysfunction, and phototransduction pathway could collectively affect ERG response (17). Similarly, the CMP-N-acetyl neuraminate biosynthesis I pathway involved sialylation of membrane proteins, regulation of synaptic transmission and ERG response (18) was also reduced and the Sirtuin signaling that is known to be protective for the retina was also inhibited (19). Additionally, activation of synaptic long-term depression and neurite outgrowth was also predicted in the ‘entrained’ retina, suggesting a possible role of active homeostatic plasticity mechanisms in response to photoreceptor degeneration.

Interestingly, scotopic ERG response seems to be rescued in the ‘freerunning’ and ‘zigzagging’ retina, despite a reduction in photoreceptor numbers; this could be either due to the involvement of homeostatic mechanisms or metabolic changes. Activation of TCA cycle pathway and downregulation of long-term synaptic depression in these behavior groups are suggestive of this change. Many recent studies have reported compensatory dendritic synapse formation in retinal bipolar cells when rod or cone photoreceptor was ablated (20). These rod or cone bipolar dendrites may extend their arbors and form new synapses to compensate for a reduced number of photoreceptor cells (21). A recent study also reported functional compensation in the retina after ablation of 50% of rods, without any observable anatomical changes in the bipolar cells (22). We believe these homeostatic mechanisms may have also contributed to rescuing photopic ERG in the entrained mice in spite of a significant reduction in cone photoreceptor numbers.

How these findings of a change in visual function will translate into humans would be an important area of research because chronically disrupted light/dark cycle could put individuals at the risk of developing more severe ocular conditions such as diabetic retinopathy (DR) and age-related macular degeneration (AMD). In a longitudinal study involving humans, retinal thinning was observed preceding the development of diabetic retinopathy (23). Also, we previously reported that the retinal clock’s genetic disruption by conditional deletion of Bmal1 accentuated the response to eye’s microvascular injury (11) and mutation in clock gene Per2 recapitulated the phenotype of DR (12). These data suggest that the retinal phenotype observed in this study could be preceding possible vascular pathology. Therefore, we speculate that chronic CRD could lead to photoreceptor degeneration and ophthalmic diseases.

In addition to visual dysfunction in response to CRD, in this study, we report different entrainment behaviors in otherwise healthy, C57BL/6J mice in response to the L10:D10 cycle. Previous studies show that most mice entrain under similar conditions and exhibit a shortened freerunning period when exposed to constant darkness (24–26). However, some reports suggest that C57BL/6J mice do not entrain to these conditions and no specific activity patterns were described (25–27). Davis and Menakar reported that mice that did not entrain to L10:D10 had a higher freerunning period (23.686 ± 0.083) than the average (23.224 ±0.044) (25). Interestingly, in our study, five out of eight ‘freerunning’ mice showed a higher than average period of 23.55±0.073. Thus, it is possible that mice in the ‘freerunning’ group had higher endogenous freerunning periods, and the observed behavior might be attributed to that.

We also observed a fascinating entrainment behavior where the mice were ‘zigzagging’ between two periods. These mice likely exhibit advancement in a circadian phase when light exposure was at the end of their dark or active phase. This phase advancement possibly continued to reduce the period length until the light exposure at the beginning of the dark/active phase delayed the activity onset. Recurrent phase advances and phase delays could lead to switching of a period every 3-5 days resulting in a ‘zigzagging’ actogram. The individual variation in the ability to undergo a phase shift could be the reason for the observed zigzagging behavior. Previous studies suggest that the inter-individual difference in the freerunning period correlates with the extent of phase shift in response to a resetting light stimulus (28). While mice used in our study were from the same genetic background, individual variations in the organization of retinal and/or SCN neuronal network could result in interindividual differences in the free-running period and entrainment behaviors. Further studies are necessary to characterize these behaviors better.

The mechanism of circadian disruption among these behavioral groups could be different. The ‘entrained’ mice showed clear activity onsets at lights off, suggesting their brain clock aligned itself to the shorter 20-hour cycle. It is plausible that their retinal clock was also aligned to a 20-hour cycle via local entrainment by light exposure (7). Adapting into a much shorter period inconsistent with the endogenous ~24 hr cycle might have caused disruptions in the circadian regulation of homeostatic processes in these mice’s retina. The ‘freerunning’ mice in our study were exposed to light and dark at all phases of their circadian cycle. Light exposure at the wrong (active) time could result in circadian disruption. Also, retinal damage caused by light is found to be greater in the subjective night and is found to be potentiated by melatonin, which usually is high at night (29, 30). Similarly, ‘zigzagging’ mice were also exposed to light at the end of their active phase. Moreover, switching between two periods by the ‘zigzagging’ mice could be affecting their circadian regulation of normal physiology. In similar experiments, mice that did not entrain their activity to L10:D10 but showed 24-hour patterns of core body temperature exhibited strong resetting of SCN clock after dissection and altered phase relationship between SCN and clocks in peripheral organs (27). This suggested L10:D10 exposure reduced the circadian network’s robustness and amplitude and altered the phasing of coupling signals.

The ramifications of different entrainment behaviors reported in this study could provide an excellent platform mimicking real-life scenarios of CRD. The ‘entrained’ behavior could possibly mimic specific rotational shirk work schedules (3). The free-running behavior likely is similar to chronic jetlag schedules with light exposure on every endogenous circadian clock phase. The zigzagging mice have a varying degree of entrainment, possibly mimicking social jetlag where people tend to delay their sleep onset later in the night on weekends and sleep early on weekdays (31).

Our data provide substantial evidence for CRD and aberrant light exposure as a causative agent for photoreceptor degeneration and visual dysfunction. We uniquely show the effect of entrainment behavior on retinal physiology. Our data has broader implications in understanding the role of CRD in the pathogenesis of ophthalmic diseases and possibly mitigating these negative impacts with lifestyle changes.

## Methods

### Animals

The C57BL/6J male (5 weeks old) mice were purchased from Jackson Laboratory. After a week of acclimatization, mice were housed individually in cages with access to running wheels in sound-attenuated and ventilated isolation cabinets (Phenome Technologies, Chicago) for ten weeks. The light schedules were either 12 hour Light: 12-hour dark (L12:D12) for control or 10 hour light: 10 hour dark (L10:D10) for circadian disruption with 300 lux light intensity during the light phase, from a cool white LED source. Food and water were provided *ad libitum*. All experiments were carried out between 10 to 14 weeks while still in L10: D10 at Zeitgeber Time (ZT) 3 to 6, where ZT 0 is the ‘lights on’ time unless otherwise specified. In cases of not entrained mice, its activity onsets were monitored for coinciding with lights off, and the experimentation was performed on the following day. All animal experimentations were carried out in accordance with the National Research Council *Guide for the Care and Use of Laboratory Animals* and the Association for Research in Vision and Ophthalmology Statement for the Use of Animals in Ophthalmic and Vision Research and approved by the Institutional Animal Care and Use Committee of Indiana University School of Medicine.

### Wheel running activity

Wheel running activity was recorded every minute for ten weeks using Actimetrics (Actimetrics, Chicago) hardware and monitored and analyzed using Clocklab (Actimetrics, Chicago). The actogram data from 15 to 70 days were used to determine the period and analyzing wheel-running activity. The period was determined either by periodogram analysis or by fitting lines on 3 to 4 consecutive onsets on the actograms (T20F and T20Z). Wheel running activity counts were determined by averaging the total activity during light and dark phases and were expressed for 24 hrs to make it comparable between groups.

### Electroretinography

Electroretinogram recordings were performed with an LKC Technologies UTAS system (LKC Technologies, Inc, Gaithersburg, MD, USA) under dark and light-adapted conditions. Before testing, all mice were dark-adapted for 24 hours to nullify the effect of light history (32), and the measurements were made between Zeitgeber times 10 and 13, the reported peak time for ERG response (33). The mice were anesthetized with an i.p. Injection of ketamine (100 mg/Kg) and Xylazine (5 mg/Kg). Topical 0.5 % proparacaine HCl eye drops were applied, and the pupils were dilated by topical application of 1% tropicamide and 2.5% phenylephrine (Alcon laboratories). The eyes were kept moist using 2.5 % hypromellose ophthalmic demulcent solution (Akron). The core body temperature was maintained using a heating pad at 37.0 C. Ground needle electrode was placed on the base of the tail and the reference electrode was placed subdermally between the eyes. The gold loop electrodes (LKC Technologies, Inc, Gaithersburg, MD, USA) placed over the cornea were used for recording ERG response. The stimulus flashes were presented in a UTAS ganzfeld (LKC Technologies). For photopic ERG recordings the mice were light adapted for 10 min inside the ganzfeld prior to testing. The values for a-wave and b-wave amplitudes, their implicit times and average oscillatory potential amplitude were obtained from an inbuilt analysis tool by LKC Technologies.

### Optomotor response recording

Quantification of mice spatial vision was performed by detecting the spatial frequency threshold of optomotor response behavior using OptoMotry device (CerebralMechanics, Inc.) as described earlier (34). Tracking of head movements in response to rotating sine-wave gratings (100% contrast) was recorded in free-moving mice. Spatial frequency was systematically increased in the staircase method until the animal did not respond, and the highest spatial frequency the animal could track was identified as the threshold. The threshold obtained for each eye was reported.

### Optical Coherence Tomography

The mice were anesthetized using a ketamine cocktail and pupils were dilated and kept moist as described for electroretinogram. SD-OCT images were obtained using the Bioptigen SD-OCT system. The optic nerve head was located manually and was used to center the scan. Retinal thickness was measured using manual calipers, 2 each at 0.35mm and 0.45mm away from the optic nerve head on each B-scan’s nasal and temporal axis. Three B scans from each eye were used for measurement, and the average of all the calipers of each eye is reported.

### Immunohistochemistry

Eyes for immunohistochemistry were isolated between ZT-5 to ZT-8 and fixed in paraformaldehyde (4%) in PBS overnight at 4° C. The eyes were immersed in 30% ethanol (v/v) in PBS and paraffin-embedded. Paraffin sections (5 µm) were cleared by 2 × 5 min incubation in xylene before rehydration through a graded series of ethanol [100%, 95% and 80 % (v/v) in PBS]. Epitope retrieval was performed by incubating in pre-warmed citrate buffer (pH 6), overnight at 56° C. The sections were washed in 0.3% Triton X-100 in PBS and blocked with 10% goat serum for 2 hrs at room temperature. Sections were incubated overnight at 4° C with primary antibodies diluted in 10% goat serum, washed in PBS, followed by incubation with secondary antibodies for 2hrs at room temperature. After washing in PBS, slides were mounted with Vectasheild containing DAPI for nuclei staining. Retinal sections were imaged with a confocal microscope (Zeiss), and images were analyzed with a Zeiss image processing station. Sections with optic nerve head were used for immunohistochemistry and imaging was done between 0.35 mm to 0.45 mm from the optic nerve head.

### Proteomics

The total proteins from the isolated retina were extracted using RIPA buffer and quantitated using BCA assay. The solubilized proteins were precipitated using Tricarboxylic acid and dissolved in 8M Urea in 100 mM Tris.HCl. An equal amount of protein starting material from each sample was reduced and alkylated with Tris (2-carboxyethyl) phosphine and chloroacetamide and digested with trypsin overnight. Samples were labeled with tandem mass tag (TMT) reagents (Thermo Fisher Scientific). The labeled sample was fractionated using the Pierce High pH Reversed-Phase Peptide Fractionation Kit (Thermo Fisher Scientific) and subject to liquid chromatography–MS/MS analysis using Orbitrap Fusion Lumos mass spectrometer (Thermo Scientific) coupled to an EASY-nLC HPLC system (Thermo Scientific). Data were acquired with the full MS acquisition with an Orbitrap resolution of 60,000, and MS/MS analysis was performed at a resolution of 50,000 and with a collision energy of 36. MS/MS database search was performed against *in silico* tryptic digest of mouse proteins UniProt FASTA database using SEQUEST HT within Proteome Discoverer 2.2 (PD 2.2, Thermo Fisher Scientific) with precursor mass tolerance of 10 ppm and fragment mass tolerance of 0.02 Da. Percolator False Discovery Rate was set to 1% and Peptide abundance values were normalized in Proteome Discover 2.2 (Thermo Scientific). The experiments were performed in two separate sets consisting of (1) E versus C and (2) F and Z versus C. Proteins that had significant changes in abundance in each behavioral group with respect to control were identified.

Ingenuity Pathway Analysis (IPA) (QIAGEN Inc., https://www.qiagenbioinformatics.com/products/ingenuity-pathway-analysis) was used to identify pathways associated with differentially expressed proteins. The complete dataset was uploaded to IPA. A threshold filter of p=0.05 was applied, and only experimentally observed interactions were selected for analysis. The significance in the canonical pathway function was defined by a P-value calculated using Fischer’s exact test that determines if the probability of association between proteins in the dataset and in the pathway is due to chance. Prediction activation scores (z-score) is also used as a statistical measure of the match between an expected relationship direction in a pathway and the observed protein expression. Diseases and conditions function was used to predict associated diseases of the brain and eye. Comparative analysis was done on the z-scores pathways as well as diseases and conditions to compare between entrainment groups.

### Statistical Analysis

The data were analyzed either using Welch one-way ANOVA followed by Dunnet’s test to account for heteroscedasticity or two-way ANOVA followed by post hoc multiple comparisons with Tukey’s test using GraphPad Prism version 9.0.0 for Windows, GraphPad Software, San Diego, California, USA, www.graphpad.com. Data are expressed as mean + SEM. The Welch F statistic (W) or Fisher F statistic and p values are reported in the result section. The significance level was set at p ≤ 0.05.

## Acknowledgments

This research is supported by funding support from the National Institute of Health (NIH)- National Eye Institute (NEI) grant R01EY02777903 to AB.

---

**Fig.S1. L10:D10 ‘Entained’ behavior actograms.** Double plotted actograms and their corresponding Lomb Scargle periodogram of an individual mouse showing ‘Entrained’ behavior under L10:D10 conditions (A-F).

**Fig.S2. L10:D10 ‘Freerunning’ behavior actograms.** Double plotted actograms and their corresponding Lomb Scargle periodogram of an individual mouse showing ‘Freerunning’ behavior under L10:D10 conditions (A-H). The grey area in the actogram depicts when activity was not recorded.

**Fig.S3. L10:D10 ‘Zigzagging’ behavior actograms.** Double plotted actograms and their corresponding Lomb Scargle periodogram of ‘Zigzagging’ behavior of L10:D10 mice (A-H). The grey area in the actogram depicts when activity was not recorded.

**Fig. S4. Peak latencies of ERG responses of L10:D10 mice.** Peak latencies of scotopic a-wave (A) and b-wave (B), and photopic a-wave (C) and b-wave (D) of control, ‘entrained’, ‘freerunning’ and ‘zigzagging’ mice. N=17 (L12:D12), 6 (L10:D10 E), 8 (L10:D10 F), 8(L10:D10 Z). Shown are mean + SEM.

**Fig. S5. Comparative analysis of predicted canonical pathways and diseases based on the retinal proteomic changes in L10:D10 mice.** Heat map comparing Z-scores of the top thirty predicted canonical pathways (A) in the retina of ‘entrained’, ‘freerunning’ and ‘zigzagging’ mice. Heat map comparing the Z-scores of the top thirty predicted diseases and conditions (B), manually filtered for those affecting eye and brain in the retina of ‘entrained’, ‘freerunning’ and ‘zigzagging’ mice. Positive z-score indicates activation (orange) and negative z-score indicate inhibition (blue).

