## Supplementary figures and images for "Circadian Rhythm Disruption Results in Visual Dysfunction"

### supplemental figures

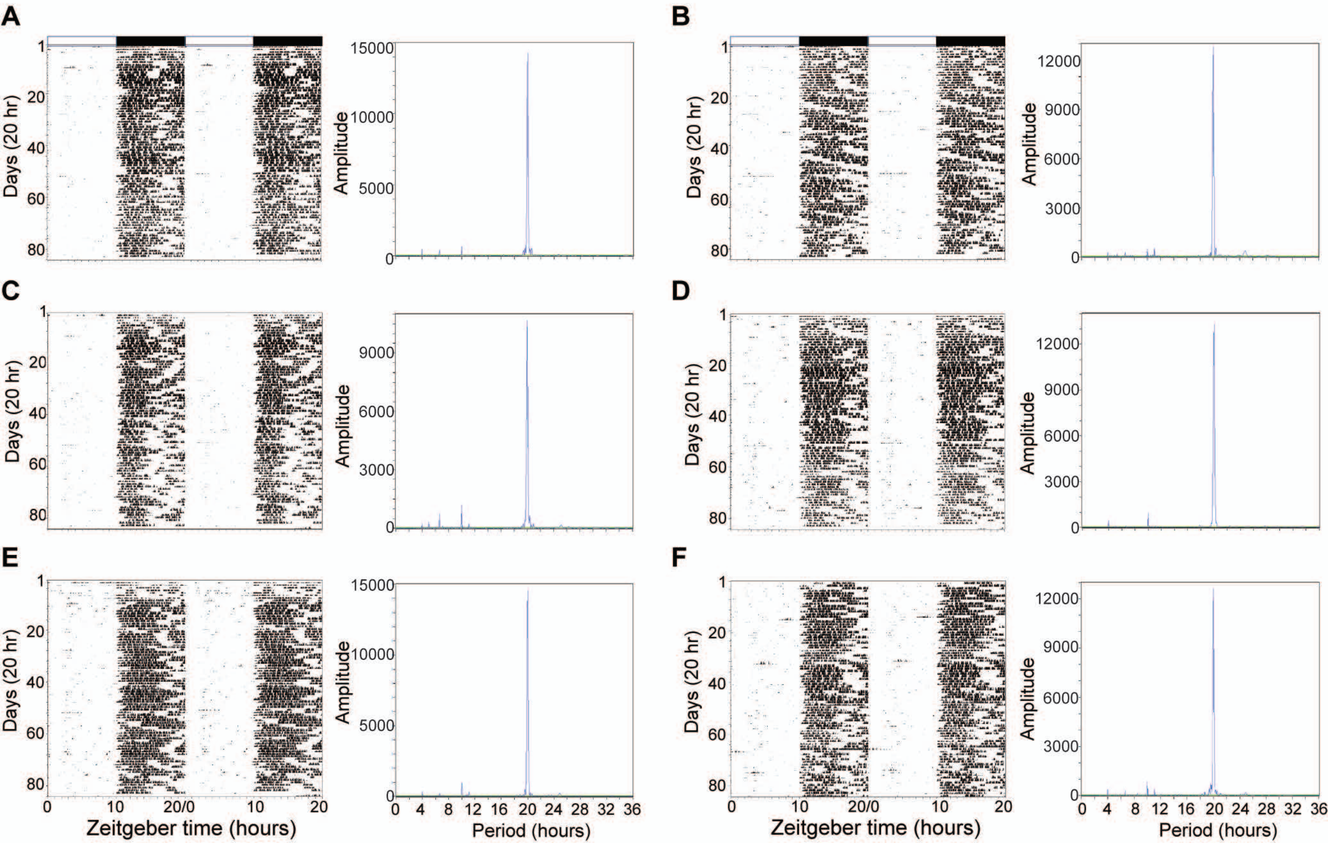

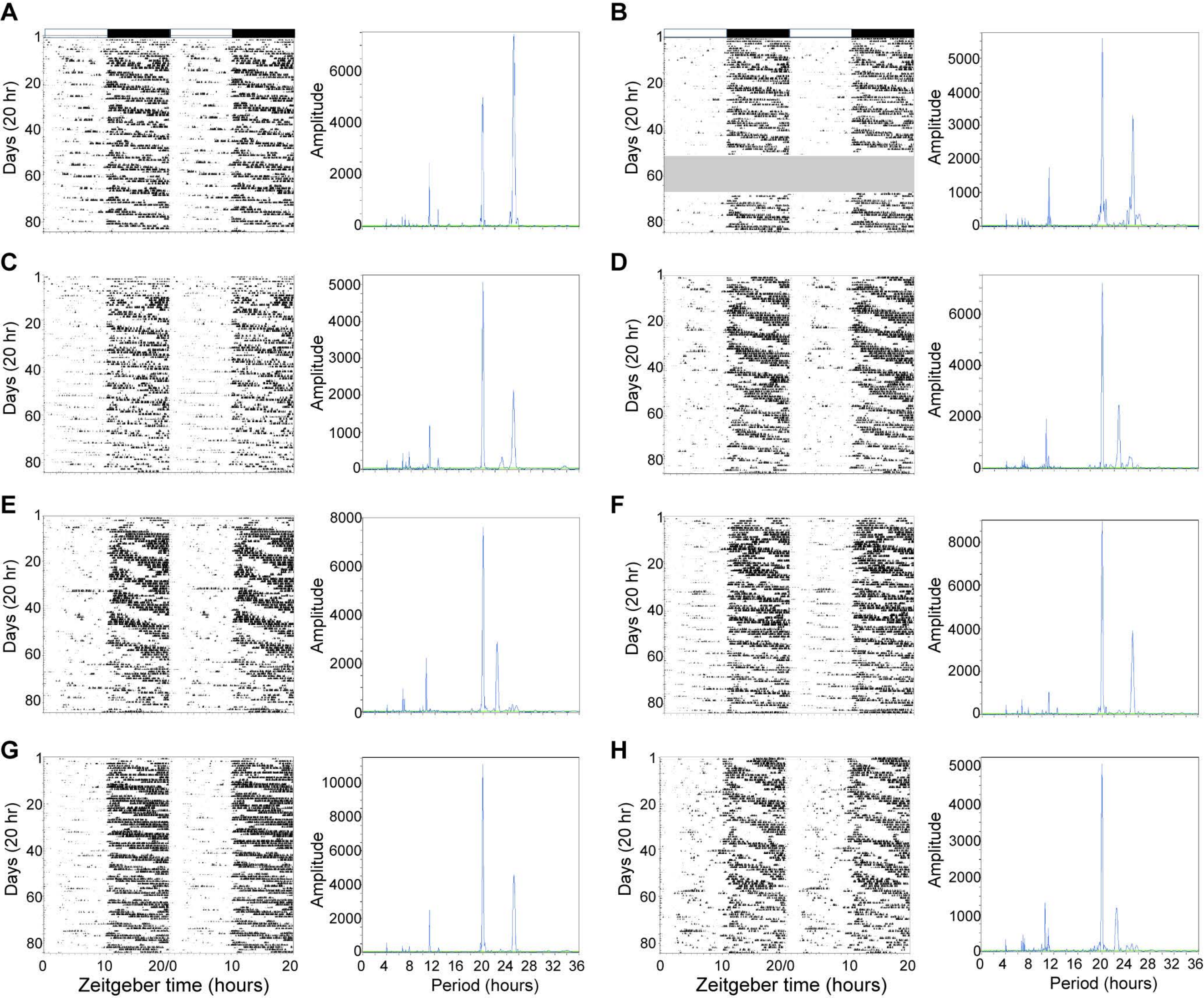

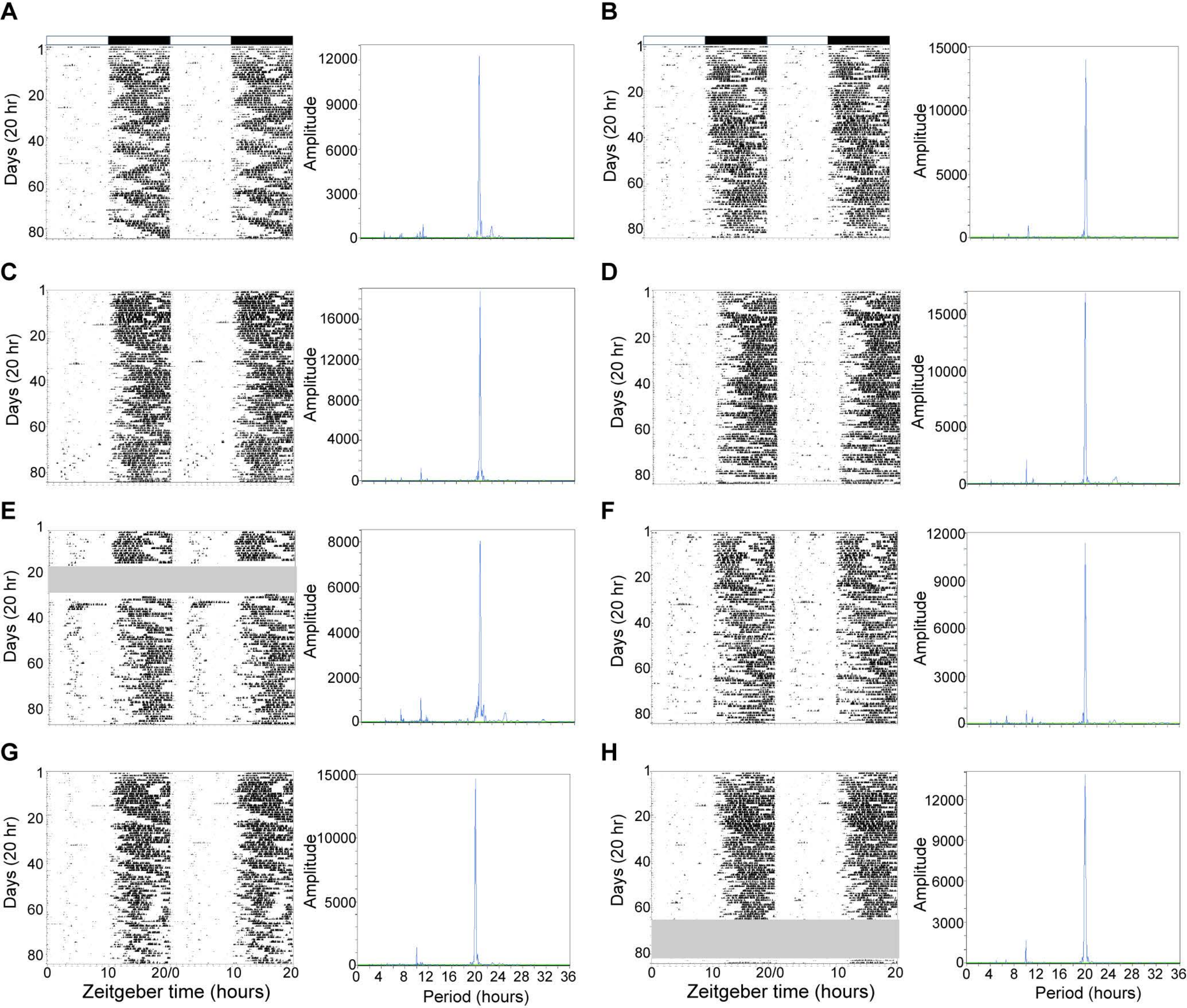

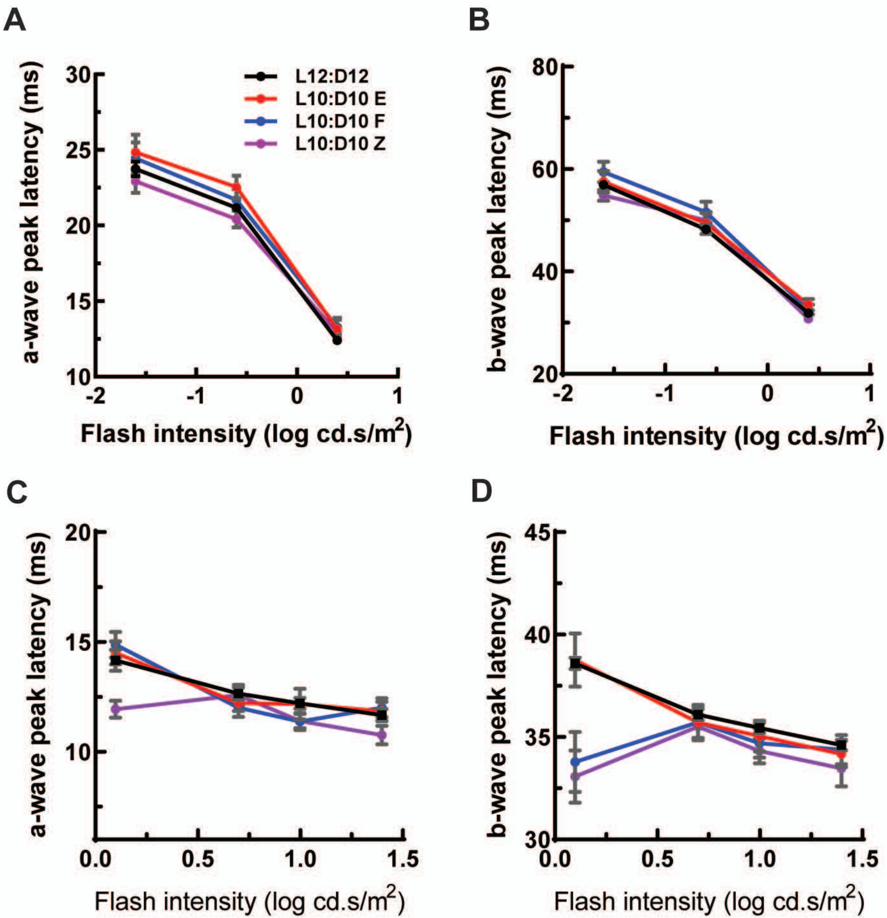

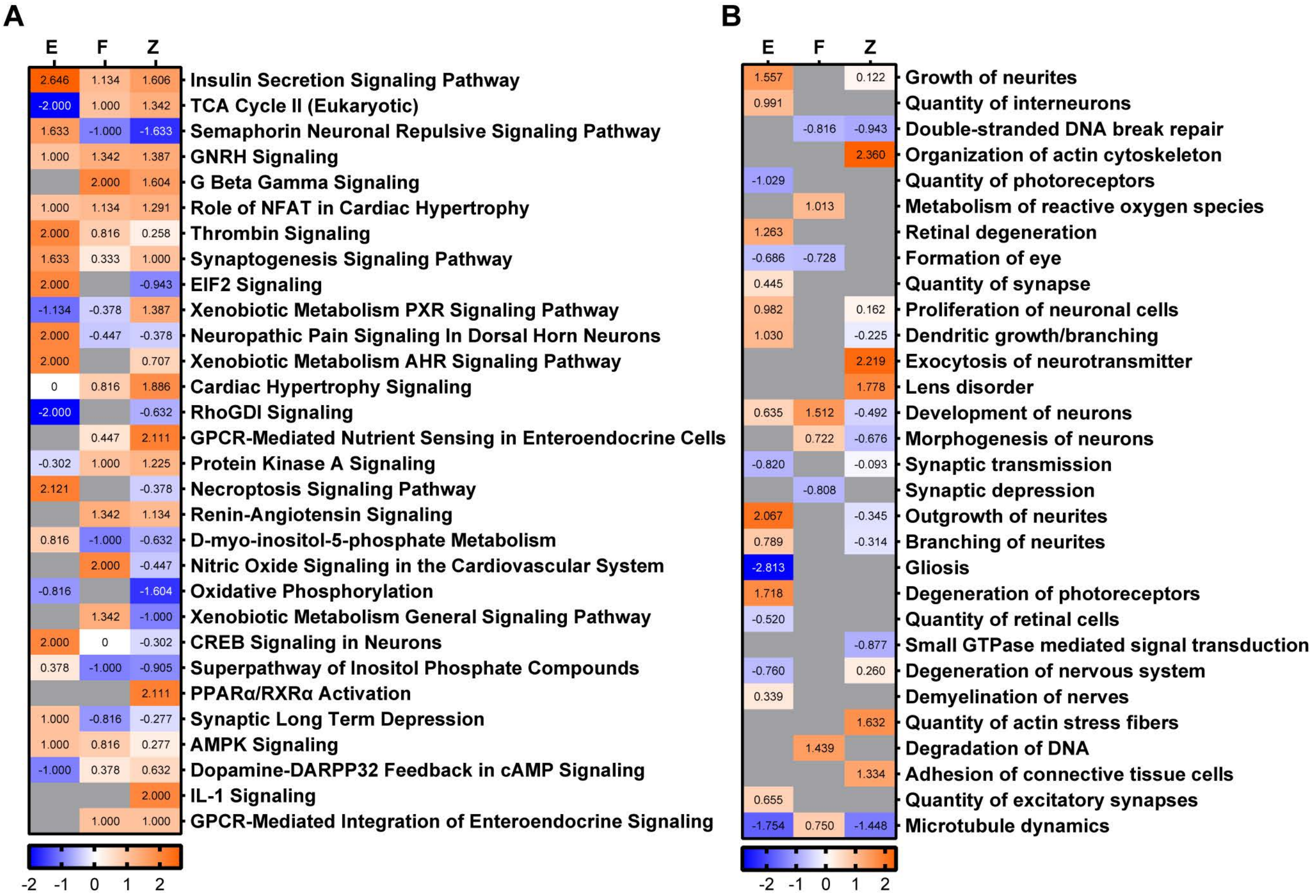
